# Distinct Microcircuit Response to Comparable Input from a Full and Partial Projection Neuron Population

**DOI:** 10.1101/2020.05.01.072629

**Authors:** Gabriel F. Colton, Aaron P. Cook, Michael P. Nusbaum

**Author notes:** G.F. Colton and A.P. Cook contributed equally to this work. 10 Linden Circle, Somerville, MA 02143, USA. To whom correspondence should be addressed: Michael P. Nusbaum, PhD, Dept. of Neuroscience, 211 Clinical Research Bldg., 415 Curie Blvd., Perelman School of Medicine, Univ. of Pennsylvania, Philadelphia, PA 19104. Author Contacts Gabriel F. Colton-, Aaron P. Cook.

## Abstract

Neuronal inputs to microcircuits are often present as multiple copies of apparently equivalent neurons. Thus far, however, little is known regarding the relative influence on microcircuit output of activating all or only some copies of such an input. We are examining this issue in the crab (*Cancer borealis*) stomatogastric ganglion, where the gastric mill (chewing) microcircuit is activated by MCN1, a paired modulatory projection neuron. Both MCN1s contain the same cotransmitters, influence the same gastric mill circuit neurons, can drive the biphasic gastric mill rhythm, and are co-activated by all identified MCN1-activating pathways. Here, we determine whether the gastric mill circuit response is equivalent when stimulating one or both MCN1s under conditions where the pair are matched to collectively fire at the same overall rate and pattern as single MCN1 stimulation. The dual MCN1 stimulations elicited more consistently coordinated rhythms, and these rhythms exhibited longer phases and cycle periods. These different outcomes from single and dual MCN1 stimulation may have resulted from the relatively modest, and equivalent, firing rate of the gastric mill neuron LG during each matched set of stimulations. The LG neuron-mediated, ionotropic inhibition of the MCN1 axon terminals is the trigger for the transition from the retraction to protraction phase. This LG neuron influence on MCN1 was more effective during the dual stimulations, where each MCN1 firing rate was half that occurring during the matched single stimulations. Thus, equivalent individual- and co-activation of a class of modulatory projection neurons will not necessarily drive equivalent microcircuit output.

**Summary Statement:** Co-stimulating both copies of an identified modulatory projection neuron at the same collective firing rate used for single copy stimulation results in distinct microcircuit output.

## INTRODUCTION

Projection neurons that regulate circuit activity commonly occur as populations of apparently equivalent, coactivated copies (Rosen et al., 1991; Blitz et al., 1999; Brodfuehrer and Thorogood, 2001; Hägglund et al., 2010; Betley et al., 2013; Bidaye et al., 2014; Gunaydin et al., 2014; Daghfous et al., 2016; Qiu et al., 2016; Li et al., 2017; Fino et al., 2018; Li and Soffe, 2019; Ruder and Arber, 2019). While the consequences of their coactivation for circuit output and/or behavior are established in many systems, there appears to be no systematic comparison of a circuit response to co-activating an entire projection neuron population versus a defined subset of it. Such comparisons would help elucidate how these systems operate, and could inform whether continued normal behavioral function would likely occur after some fraction of the population was compromised due to injury or disease (Fink and Cafferty, 2016).

The circuit response to activating all or some members of a projection neuron population might be equivalent, particularly when those neurons act at least partly via peptide cotransmitters, because neurally-released peptides often diffuse broadly and have long-lasting effects on their targets (Marder, 2012; van den Pol, 2012; Nusbaum et al., 2017; Svensson et al., 2019). Alternatively, the circuit response might be skewed by the number of active projection neurons and their firing rates, for example because neuropeptide release often has a higher firing rate threshold than small molecule co-transmitters and its release can be a non-linear function of firing rate (Cazalis et al., 1985; Peng and Horn, 1991; Whim and Lloyd, 1994; Vilim et al., 1996, 2000; Arrigoni and Saper, 2014; Nusbaum et al., 2017; Blitz et al., 2019), and modulatory actions can be concentration-specific (Flamm and Harris-Warrick, 1986; Saideman et al., 2006; Fort et al., 2007; Dickinson et al., 2015; Blitz et al., 2019). In some cases, an apparently equivalent projection neuron population was subsequently determined to diverge and influence overlapping or distinct sets of neurons (Lammel et al., 2011; Betley et al., 2013; Luo et al., 2018).

Manipulating the activity of projection neuron populations in a controlled manner is possible, for example by optogenetics, but it remains challenging to precisely manipulate a defined subset of them. Such precision is possible, however, in some smaller systems where identified projection neurons with a known influence on a target circuit are present as small copy numbers (Rosen et al., 1991; Frost and Katz, 1996; Blitz et al., 1999; Brodfuehrer and Thorogood, 2001; Mesce et al., 2008; Jeanne and Wilson, 2015).

One tractable system for such a study is the stomatogastric nervous system (STNS) of the crab *Cancer borealis* (Marder and Bucher, 2007; Daur et al., 2016; Nusbaum et al., 2017; Stein, 2017). This system contains two well characterized microcircuits in the stomatogastric ganglion (STG) which generate the motor patterns for chewing (gastric mill circuit) and the pumping/filtering of chewed food (pyloric circuit) in vivo and in the isolated STNS. These circuits are readily accessible in the isolated STNS because most of the twenty-six STG neurons contribute to one or both of these circuits, their somata are relatively large (diameter: ∼30 – 120 µm), and most circuit neurons occur as single copies. There are also six different identified projection neuron pairs which regulate gastric mill- and/or pyloric circuit activity (Coleman et al., 1992; Norris et al., 1994, 1996; Blitz et al., 1999; Christie et al., 2004).

The best characterized projection neuron in the *C. borealis* STNS is modulatory commissural neuron 1 (MCN1) (Coleman and Nusbaum, 1994; Hedrich et al., 2011; Nusbaum et al., 2017). Each MCN1 projects an axon from one of the paired commissural ganglia (CoGs), through the inferior oesophageal nerve (*ion*) and stomatogastric nerve (*stn*), to innervate the STG and drive the gastric mill and pyloric rhythms (Coleman et al, 1992; Coleman and Nusbaum, 1994; Bartos and Nusbaum, 1997). MCN1 can be selectively driven by extracellular *ion* stimulation, and both MCN1s contain the same cotransmitters, influence the same STG circuit neurons, and drive the gastric mill rhythm by the same mechanism (Coleman and Nusbaum, 1994; Coleman et al., 1995; Bartos and Nusbaum, 1997; Bartos et al., 1999; Blitz et al., 1999; Wood et al., 2000, 2004; Beenhakker and Nusbaum, 2004; Stein et al., 2007; DeLong et al., 2009a,b; Nusbaum et al., 2017). Furthermore, despite being present in separate ganglia, both MCN1s are coactivated by all identified sensory and CNS pathways (Beenhakker et al., 2004, 2005; Blitz et al., 2004, 2008; Christie et al., 2004; Hedrich et al., 2009; Blitz and Nusbaum, 2012; White et al., 2017).

Here, we test the hypothesis that the gastric mill circuit response to dual and single MCN1 stimulation, in preparations where the CoGs are removed and the MCN1 axon in the *ion* is selectively stimulated, is comparable when the two conditions have matched firing rates and patterns. We ensure this match by co-stimulating the two MCN1s in a one-to-one alternating pattern that produces an inter-stimulus interval equivalent to that used for the matched single MCN1 stimulation. These matched stimulations did not elicit equivalent gastric mill rhythms. For example, the gastric mill rhythm was more consistently coordinated during the dual MCN1 stimulations. These different outcomes likely resulted at least partly from the gastric mill rhythm generator neuron LG (lateral gastric) firing frequency being the same within each set of matched MCN1 stimulations. The lower firing rate of each MCN1 during the dual stimulations, relative to the matched single stimulations, apparently resulted in them being more strongly inhibited by LG. These results indicate that at least some neural circuits are optimally driven by co-activating the full population of circuit-driving projection neurons.

## METHODS

### Materials and Methods

#### Animals

Male Jonah crabs (*Cancer borealis*) were obtained from commercial suppliers (Fresh Lobster, LLC; Marine Biological Laboratory; Ocean Resources, Inc.) and maintained in aerated, filtered artificial seawater at 10–12° C. Animals were cold anesthetized by packing in ice for at least 30 min before dissection. The foregut was then removed from the animal, after which the STNS was dissected from the foregut in physiological saline at 4°C. The dorsal connective tissue sheath of the STG was removed immediately prior to recording to facilitate access for intrasomatic recordings.

#### Solutions

*C. borealis* physiological saline contained the following (in mM): 440 NaCl, 26 MgCl_2_, 13 CaCl_2_, 11 KCl, 10 Trizma base, 5 maleic acid, and 5 glucose, pH 7.4 – 7.6. All preparations were superfused continuously with *C. borealis* saline (8 –12° C).

#### Electrophysiology

Electrophysiology experiments were performed using standard techniques for this system (Beenhakker and Nusbaum, 2004). The isolated STNS (Fig. 1A) was pinned down in a silicone elastomer-lined (Sylgard 184: KR Anderson) Petri dish. Each extracellular nerve recording resulted from a pair of stainless steel wire electrodes (reference and recording) whose ends were pressed into the Sylgard-coated dish. A differential AC amplifier (model 1700: A-M Systems) amplified the voltage difference between the reference wire, placed in the bath, and the recording wire, placed near an individual nerve and isolated from the bath by petroleum jelly (Vaseline; Lab Safety Supply). This signal was then further amplified and filtered (model 410 amplifier: Brownlee Precision). Extracellular nerve stimulation was accomplished by placing the pair of wires used to record nerve activity into a stimulus isolation unit (SIU 5: AstroNova Inc.) that was connected to a stimulator (model S88: AstroNova Inc.).

**Figure 1.**
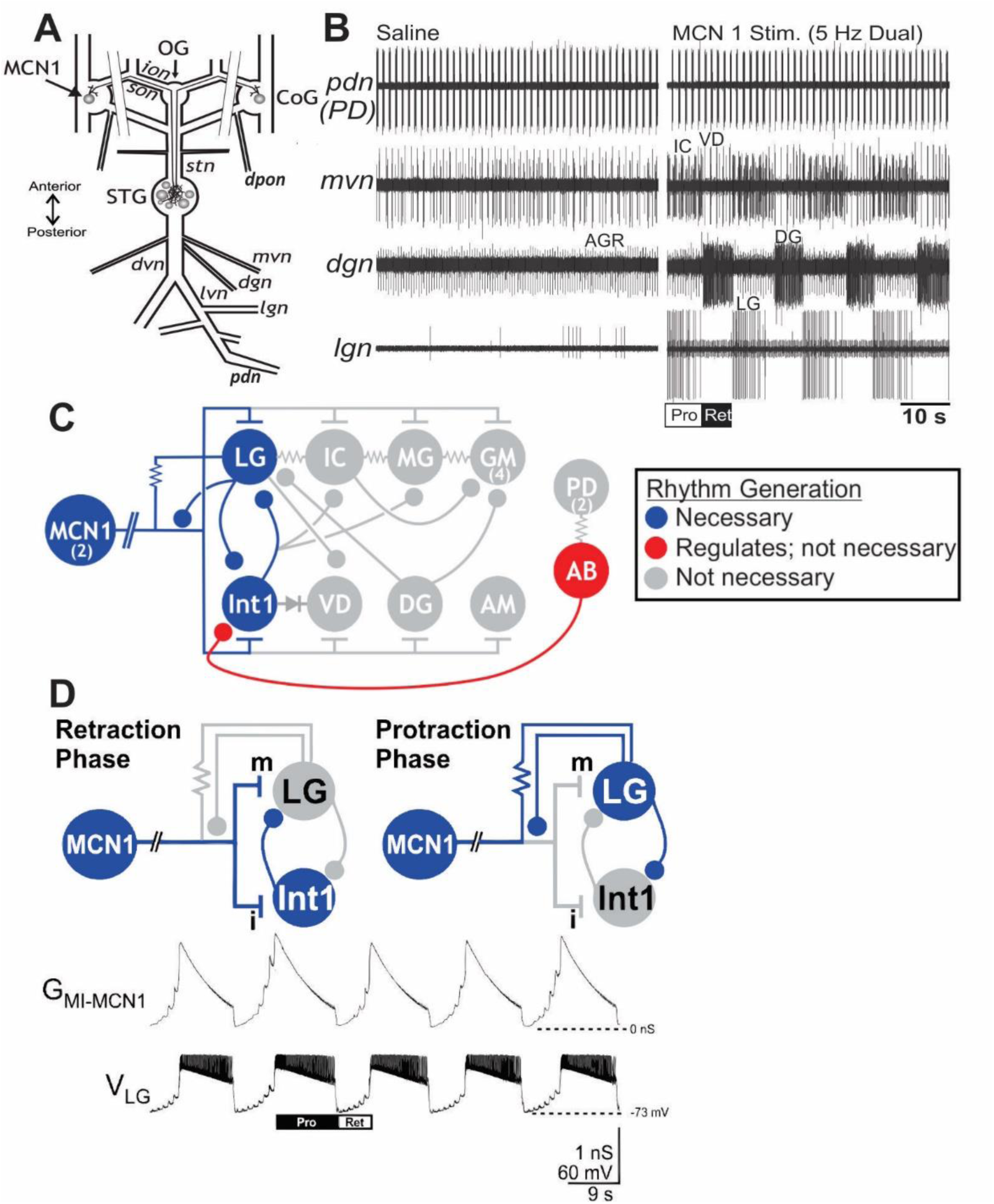
The projection neuron MCN1 drives the gastric mill rhythm in the isolated stomatogastric nervous system of the crab *Cancer borealis*. (A) STNS schematic, with the parallel lines crossing the *ions* and *sons* representing the nerve transections used to remove the CoGs. (B) Selective MCN1 (*ion*) stimulation drives the gastric mill rhythm. Before stimulation (Saline), there was an ongoing pyloric rhythm (*pdn, mvn*) but no gastric mill rhythm (*dgn, lgn*). Small unit in *mvn*: GM neuron; small continuously present units in the *lgn* are stimulation artifacts. AGR (anterior gastric receptor) is a sensory neuron that only influences the gastric mill rhythm via its actions in the CoGs (Simmers and Moulins, 1988; Norris et al., 1994; Smarandache and Stein, 2007; Hedrich et al., 2009). Pro, protraction; Ret, retraction. (C) Schematic of the MCN1-driven gastric mill circuit. Top row, protractor neurons; bottom row-retractor neurons. Labels: Filled circles, inhibition; T-bars, excitation; Resistors, non-rectifying electrical coupling; Diode: rectified electrical coupling; Double slashed lines on MCN1 axon: space break. Modified from: Saideman et al. (2007). (D) (top) Schematic of the phase specific operation of the MCN1-gastric mill rhythm generator circuit. Labels: blue represents active neurons and synapses; grey represents inactive neurons and synapses. (bottom) Computational model output of the MCN1-gastric mill rhythm activity in the LG neuron plus the associated, rhythmic waxing and waning of MCN1-activated g_MI_ in LG (g_MI-MCN1_). Modified From: DeLong et al. (2009a).

Intrasomatic recordings were made with sharp glass microelectrodes (8 – 20 MΩ) filled with potassium acetate (KAcetate: 4 M) plus KCl (20 mM), or KCl alone (1 M). Intracellular signals were amplified using Axoclamp 2B amplifiers (Molecular Devices), and then further amplified and filtered (Brownlee model 410 amplifier). Current injections were performed in single-electrode discontinuous current-clamp (DCC) mode with sampling rates between 2 – 3 kHz. To improve visibility for intracellular recording, the STG was dorsally desheathed and viewed with light transmitted through a dark-field condenser (Nikon). STG neurons were identified on the basis of their axonal projections, activity patterns, and interactions with other STG neurons (Weimann et al., 1991; Blitz et al., 2008).

Each *ion* was stimulated (duration per stimulus: 1 ms) to selectively activate either MCN1_Left_ (MCN1_L_) or MCN1_Right_ (MCN1_R_) (Bartos and Nusbaum, 1997). The left and right CoG, and hence MCN1_L_ and MCN1_R_, were defined based on the isolated STNS being pinned with the dorsal side facing up. There is only one other CoG projection neuron (MCN5) that innervates the STG which has an axon in the *ion*, and it has a higher stimulus threshold than MCN1 as well as having no direct influence on LG (Norris et al., 1996; Blitz et al., 2019). Each suprathreshold *ion* stimulus elicited a single MCN1 action potential, as established by recording the resulting unitary, electrical excitatory postsynaptic potential (eEPSP) in the LG neuron (Coleman et al., 1995). A tonic *ion* stimulation pattern (i.e. constant inter-stimulus intervals) was used in all these experiments.

To determine the relative influence of matched dual vs. single MCN1 stimulation on the gastric mill rhythm, we used a range of MCN1 firing rates that spanned its activity range in response to input pathway stimulation. Specifically, we stimulated each MCN1 separately, at 10 Hz, 20 Hz or 30 Hz using a tonic stimulation pattern. We compared these single stimulations to matched dual MCN1 stimulation during which the two MCN1s were each stimulated at 50% the firing rate of the associated single stimulations (e.g. 5 Hz + 5 Hz compared with 10 Hz). During dual stimulation, each MCN1 was activated to fire a single action potential alternately, such that their combined firing rate and pattern was the same as the matched single stimulation.

In some experiments where the activity of the retraction phase neuron DG persisted through the protraction phase of the gastric mill rhythm, the ability of the protractor neuron LG to regulate DG activity was examined by injecting LG with supra-threshold depolarizing current pulses during some of its bursts to increase its within-burst firing rate. In these same experiments, we also determined the impact of these LG current injections on the pyloric cycle period (see below). The LG current injections (+1 – +4 nA) were done in DCC, using brief depolarizing pulses (pulse duration: 30 – 50 ms) that each elicited a single action potential. Each within-burst current injection train was turned on and off manually, and was timed to occur during a normal LG burst for a duration similar to the LG burst duration during that particular gastric mill rhythm. These manipulations were performed in a subset of experiments during dual 10 Hz- and single 20 Hz MCN1 stimulated gastric mill rhythms. Two of these LG current injections, separated by 4-8 control LG bursts, were performed during each gastric mill rhythm.

In control experiments, the mechanosensory ventral cardiac neurons (VCNs) were activated by stimulating the dorsal posterior oesophageal nerve (*dpon*: duration per stimulus: 1 ms), either uni- or bilaterally, in preparations where the superior oesophageal nerves (*sons*) were bisected between the *dpon* and the stomatogastric nerve (*stn*). The VCNs innervate the ipsilateral CoG via the *dpon* and *son*, and their stimulation triggers a version of the gastric mill rhythm by eliciting a long-lasting activation of the CoG projection neurons MCN1 and CPN2 (commissural projection neuron 2) (Beenhakker et al., 2004; Beenhakker and Nusbaum, 2004). Bisecting the *sons*, through which the CPN2 axon projects to the STG, prevents CPN2 activity from influencing the STG, enabling VCN stimulation to influence the gastric mill circuit exclusively via its activation of MCN1 (Norris et al. 1994; Beenhakker and Nusbaum, 2004). These experiments included both bilateral (n=2) and unilateral VCN stimulations (n=3). In each experiment, the MCN1 firing rate resulting from VCN stimulation was determined on the stimulated side(s) and subsequently used as the ipsilateral *ion* stimulation frequency, to compare the gastric mill rhythm response to direct (*ion* stimulation) and indirect (*dpon* stimulation) activation of MCN1. All *ion* stimulation experiments were performed from Sept, 2013 – March, 2015. The *dpon* vs. *ion* stimulation experiments were performed during Feb. – March, 2017.

#### Data analysis

Data were collected in parallel onto a chart recorder (AstroNova Everest model) and computer. Acquisition onto computer (sampling rate ∼5 kHz) used the Spike2 data acquisition and analysis system (Cambridge Electronic Design). Some analyses, including cycle period, burst durations, duty cycle, number of action potentials per burst, and intraburst firing frequency were conducted on the digitized data using a custom-written Spike2 program (‘The Crab Analyzer’). To facilitate data analysis and improve clarity in some figures, a raw extracellular recording (e.g. *dgn*) was duplicated with the stimulation artifacts digitally subtracted, reducing their amplitude or eliminating them. This is indicated where appropriate in each figure legend, and is illustrated in Figure 2B. A custom written script in Spike2 was used to digitally subtract the artifacts after manually inspecting the trace to verify that only that particular unit was selected for subtraction (Blitz and Nusbaum, 2008).

**Figure 2.**
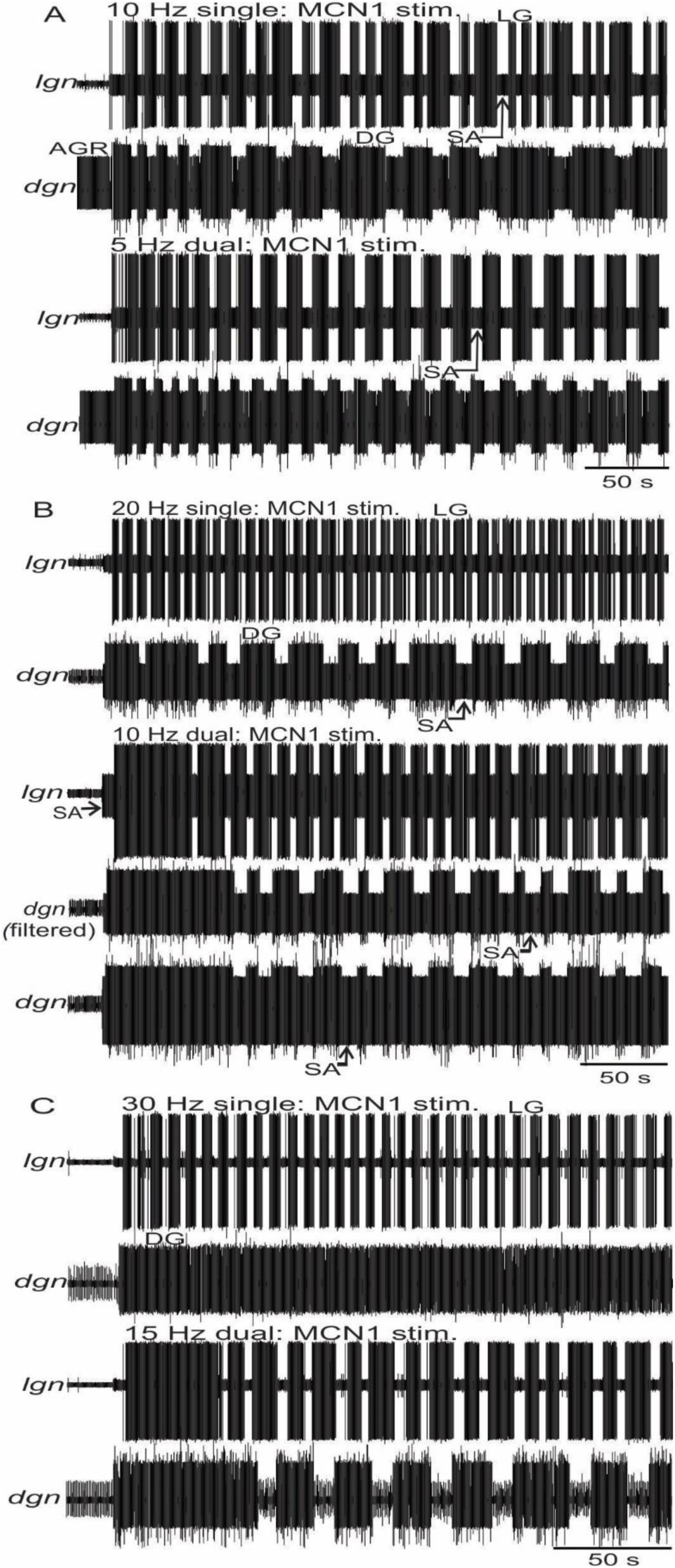
Example recordings of gastric mill rhythms in response to firing rate- and pattern matched dual and single MCN1 stimulations at a compressed time scale. Each rhythm is represented by extracellular recordings of the protractor LG (*lgn*) and retractor DG (*dgn*) neurons. Thickened baseline represents stimulation artifacts (SA). Each matched pair comes from the same experiment, but the different panels come from different experiments. Where noted in this and succeeding figures, nerve recordings were filtered to digitally reduce stimulus artifact size to better separate them from the recorded action potentials. (A) 5 Hz dual vs 10 Hz single MCN1 stimulation. (B) 10 Hz dual vs. 20 Hz single MCN1 stimulation, including *dgn* without vs. with filtering in the ‘10 Hz dual’ panel. (C) 15 Hz dual vs. 30 Hz single MCN1 stimulation. Filtered *dgn*: all recordings except 20 Hz single stimulation.

Unless otherwise stated, each data point in a dataset was derived by determining the mean for the analyzed parameter from 8-20 consecutive gastric mill cycles. One gastric mill cycle was defined as extending from the onset of consecutive LG neuron action potential bursts (Beenhakker and Nusbaum, 2004; Wood et al., 2004). Thus, the gastric mill cycle period was measured as the duration between the onset of two successive LG neuron bursts. The protractor phase was measured as the LG burst duration, whereas the retractor phase was measured as the LG interburst duration. A gastric mill rhythm-timed burst duration was defined as the duration between the onset of the first and last action potential within an impulse burst, during which no interspike interval was longer than 2 s (approximately twice the pyloric cycle period during the gastric mill rhythm and no more than half the duration of each gastric mill phase) (Beenhakker et al., 2004). The intraburst firing rate of a neuron was defined as the number of action potentials within a burst minus one, divided by the burst duration. The pyloric cycle period was determined as the duration between the onset of successive bursts in the pyloric dilator (PD) neuron (Fig. 1B). The PD neuron is a pyloric pacemaker neuron component (Marder and Bucher, 2007; Selverston and Miller, 1980).

To evaluate the influence of the LG neuron depolarizing current injections on the pyloric cycle period, the longest pyloric cycle that occurred after the start of the LG burst was selected for analysis during (a) each LG burst that received depolarizing current pulses, and (b) the two preceding LG bursts. The first pyloric cycle after LG burst onset was always the slowest one (i.e. longest cycle period) during the control LG bursts and was the slowest during 77% of the depolarized LG bursts, likely reflecting the fact that the LG instantaneous firing rate was often highest at the start of its burst. To compare these pyloric cycle periods between the depolarized and control LG bursts, we first determined and compared the ratios of the pyloric cycle period from the (a) depolarized LG burst divided by that from the preceding control LG burst, and (b) the same preceding control LG burst divided by that from the control LG burst two prior to the depolarized burst.

Data were plotted with Excel (version 2002; Microsoft) and Matlab (version 8; MathWorks). Figures were produced using CorelDraw (version 13.0 for Windows). Statistical analyses were performed by comparing the overall mean of individual mean values for two different manipulated conditions, or control and manipulated groups, from n experiments (see Results for each n value) using Microsoft Excel, SigmaPlot 13.0 (SPSS Inc.), and Matlab. Comparisons were made to determine statistical significance using (a) Repeated Measures Analysis of Variance (RM-ANOVA), with the Holm-Sidak post-hoc test when the RM-ANOVA p value was <0.05, (b) RM-ANOVA on Ranks, with the post-hoc Tukey test or Chi-Square test, (c) paired Student’s t-test, (d) Signed Rank test, (e) Chi-Square test (different from the aforementioned post hoc Chi-Square test), (f) Fisher’s Exact test, (g) Paired t-test, (h) Wilcoxon Signed Rank test, or (i) Unpaired t-test. In all experiments, the effect of each manipulation was reversible, and there was no significant difference between the pre- and post-manipulation groups. Significant differences were determined to occur when p<0.05. Data are expressed as the mean ± standard error.

## RESULTS

The gastric mill rhythm is a two-phase motor pattern (protraction, retraction) that drives the rhythmic contraction of striated muscles in the middle (i.e. gastric mill) stomach compartment of decapod crustaceans (Heinzel, 1988; Heinzel et al., 1993; Diehl et al., 2013). These muscle contractions cause the paired lateral teeth and medial tooth within the gastric mill to rhythmically move towards (protract) and away from (retract) the midline, macerating food moved into the gastric mill from the anterior, cardiac sac stomach compartment. The chewed food is then filtered and pumped through the posterior stomach compartment, the pylorus, to enter the midgut for nutrient absorption. The rhythmic chewing pattern is generated by the gastric mill central pattern generator circuit, an episodically active microcircuit in the STG which is driven by projection neurons, including the paired MCN1s, located in the CoGs (Fig. 1A,C) (Coleman and Nusbaum, 1994; Bartos et al., 1999; Beenhakker and Nusbaum, 2004; Blitz et al., 2008, 2019). The gastric mill rhythm is episodically active, in vivo and in vitro, because the projection neurons which drive it are episodically active, requiring excitatory drive from sensory neurons and other CNS neurons (Beenhakker et al., 2004; Blitz et al., 2004, 2008; Christie et al., 2004; Hedrich et al., 2009, 2011; Diehl et al., 2013).

There are eight gastric mill circuit neuron types, including four protractor motor neurons (LG, MG, GM, IC) and four retraction phase neurons (one interneuron: Int1; three motor neurons: DG, VD, AM) (Fig. 1C) (Nusbaum et al., 2017). All but GM are present as single copies; there are four GMs. The core rhythm-generating sub-circuit during selective MCN1 activation includes the reciprocally inhibitory pair LG-Int1 plus the STG terminals of MCN1 (MCN1_STG_) (Fig. 1D) (Coleman et al., 1995; Bartos et al., 1999; DeLong et al., 2009a; Nusbaum et al., 2017).

MCN1 is a multi-transmitter neuron which uses only its peptide cotransmitter CabTRP Ia (*Cancer borealis* tachykinin-related peptide Ia) to influence LG, causing a slowly developing excitation by activating I_MI_ (<u>m</u>odulator-activated inward current), while it uses only its small molecule cotransmitter GABA to influence Int1, causing a fast excitation (Blitz et al., 1999; Wood et al., 2000; Stein et al., 2007). MCN1 also has a functionally important electrical synapse with LG which strengthens the LG burst, and it receives fast glutamatergic inhibition onto its STG terminals from LG which triggers the transition from retraction to protraction and enables the eventual transition from protraction to retraction (Fig. 1D) (Coleman and Nusbaum, 1994; Coleman et al., 1995; DeLong et al., 2009a). Here, we will use the LG neuron burst to represent the protraction phase, and the LG neuron interburst burst to represent the retraction phase (Fig. 1B).

### Distinct LG/DG neuron coordination during dual vs. single MCN1 stimulation

We examined gastric mill rhythms resulting from three sets of matched dual vs. single MCN1 stimulation (5 Hz dual vs. 10 Hz single; 10 Hz dual vs. 20 Hz single; 15 Hz dual vs. 30 Hz single). All stimulation protocols elicited the gastric mill rhythm, although each matched dual vs single stimulation differed with respect to the likelihood of driving a fully coordinated motor pattern (Figs. 2,3).

**Figure 3.**
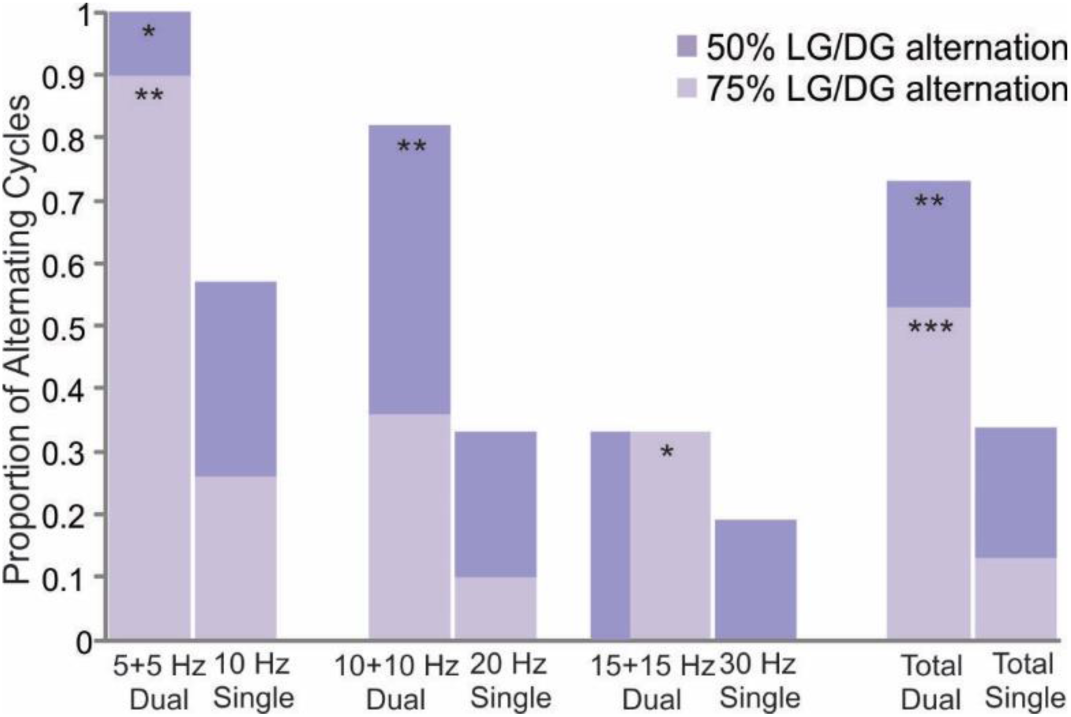
Proportion of MCN1-gastric mill rhythms in which the LG and DG neurons burst in alternation for at least 50% or 75% of the associated cycles. Bar graphs comparing the proportion of gastric mill rhythms in which LG/DG alternation occurred in at least 50% and 75% of the analyzed cycles during each set of matched dual and single MCN1 stimulations. Note that the y-axis value was the same for the 50% and 75% analyses for the 15 Hz Dual condition. Statistical analysis-Fisher’s Exact Test; Chi-Square Test on Contingency Table [total dual vs. total single]: *p<0.05; **p<0.01; ***p<0.001.

A fully coordinated gastric mill rhythm exhibits consistent alternating bursting between the protraction- and retraction phase neurons (Fig. 1B,C). Most aspects of this patterning during the MCN1-driven gastric mill rhythm result directly from the circuit synapses made by the rhythm generator neurons LG and Int1 (Fig. 1C) (Bartos et al., 1999; White and Nusbaum, 2011). Consequently, most gastric mill neuron activity is tightly linked to that of LG and/or Int1. For example, IC neuron activity is consistently limited to the protraction phase (LG burst) while VD neuron activity is limited to the retraction phase (LG interburst) (Figs. 1B, 4). In contrast, the gastric mill rhythm-timed bursting pattern of the retractor neuron DG is an indirect consequence of LG neuron activity. Specifically, the DG burst pattern results from the LG ionotropic inhibition of MCN1_STG_, which weakens or eliminates MCN1 metabotropic excitation of DG (Fig. 1C,D) (Coleman and Nusbaum, 1994). As is evident in Figure 2, in our matched MCN1 stimulation experiments DG neuron activity was not always limited to the retraction phase.

For each of the matched pairs of dual- and single MCN1 stimulation, the dual stimulations were more effective at restricting DG neuron activity to the retraction phase (Figs. 2,3). For example, setting a threshold where LG/DG bursts alternated in at least 75% of ten – twenty consecutive gastric mill rhythm cycles, the 5 Hz dual MCN1 stimulation surpassed this threshold more consistently than the matched 10 Hz single stimulation (5 Hz dual-n=9/10, 90%; 10 Hz single-n=5/19, 26%; p=0.002, Fisher’s Exact test) (Fig. 3). There was a comparable distinction at the highest matched MCN1 stimulation frequencies (15 Hz dual vs. 30 Hz single: p= 0.037, Fisher’s Exact test), although not during the matched 10 Hz dual and 20 Hz single MCN1 stimulations (p=0.148) (Fig. 3). Additionally, in eight of the ten (80%) 5 Hz dual stimulation experiments there was LG/DG burst alternation in every gastric mill rhythm cycle. Comparable full coordination for all cycles only occurred during three of eleven (27%) 10 Hz dual stimulations and none (0/9) of the 15 Hz dual stimulations. During the single MCN1 stimulations, full coordination during all cycles did not occur during any 10 Hz (n=19) or 30 Hz (n=16) stimulations, and in only one of twenty-one (5%) 20 Hz stimulations.

When threshold for the percentage of gastric mill cycles exhibiting LG/DG alternation was lowered to 50%, the two lowest dual stimulation sets were further improved (5 Hz: n=10/10, 100%; 10 Hz: n=9/11, 82%). This resulted in a significant difference for the middle matched set (10 Hz dual, n=9/11, 82%; 20 Hz single: n=5/21, 24%; p=0.003, Fisher’s Exact test), whereas the dual and single stimulations in the highest matched set became indistinguishable (p=0.63) (Fig. 3).

Collectively, only 13% of single MCN1 stimulations (n=7/56) produced consistent LG/DG burst alternation in at least 75% of cycles across all three single stimulation rates (Fig. 3). In contrast, in the same experiments, 53% (n=16/30; p<0.001, Chi Square Test on Contingency Table) of matched dual stimulations successfully drove alternating LG/DG bursting in 75% or more gastric mill rhythm cycles (Fig. 3). Lowering the threshold to 50% of cycles exhibiting LG/DG burst alternation still separated the collective single (19/56; 34%) and dual (22/30; 73%) stimulations (p=0.001, Chi Square Test on Contingency Table).

During gastric mill rhythm cycles when LG/DG burst alternation did not occur, DG neuron activity persisted through the LG burst (Figs. 2,4). When DG activity continued through the protraction phase, its firing rate was consistently reduced relative to that during retraction (10 Hz single: p<0.001, n=9; 10 Hz dual: p=0.002, n=5; 20 Hz single: p<0.001, n=10; 15 Hz dual: p=0.015, n=4; 30 Hz single: *p=0.008, n=8; Paired t-test, except *Signed Rank test). This DG firing rate reduction during protraction was greater during the dual 10 Hz and dual 15 Hz MCN1 stimulations than during their matched single stimulations (10 Hz dual vs. 20 Hz single: n=5, p=0.027; 15 Hz dual vs. 30 Hz single: n=4, p=0.037, Paired t-test). The larger DG firing rate reduction during protraction when both MCN1s were co-stimulated occurred despite the fact that the retraction phase firing rate of DG was equivalent during the matched dual and single stimulations (10 Hz dual, 12.5 ± 0.6 Hz, n=5; 20 Hz single, 12.0 ± 0.5 Hz, n=8, p=0.83; 15 Hz dual, 14.8 ± 0.8 Hz, n=4; 30 Hz single, 13.4 ± 0.3 Hz, n=8, p=0.35; Unpaired t-test).

**Figure 4.**
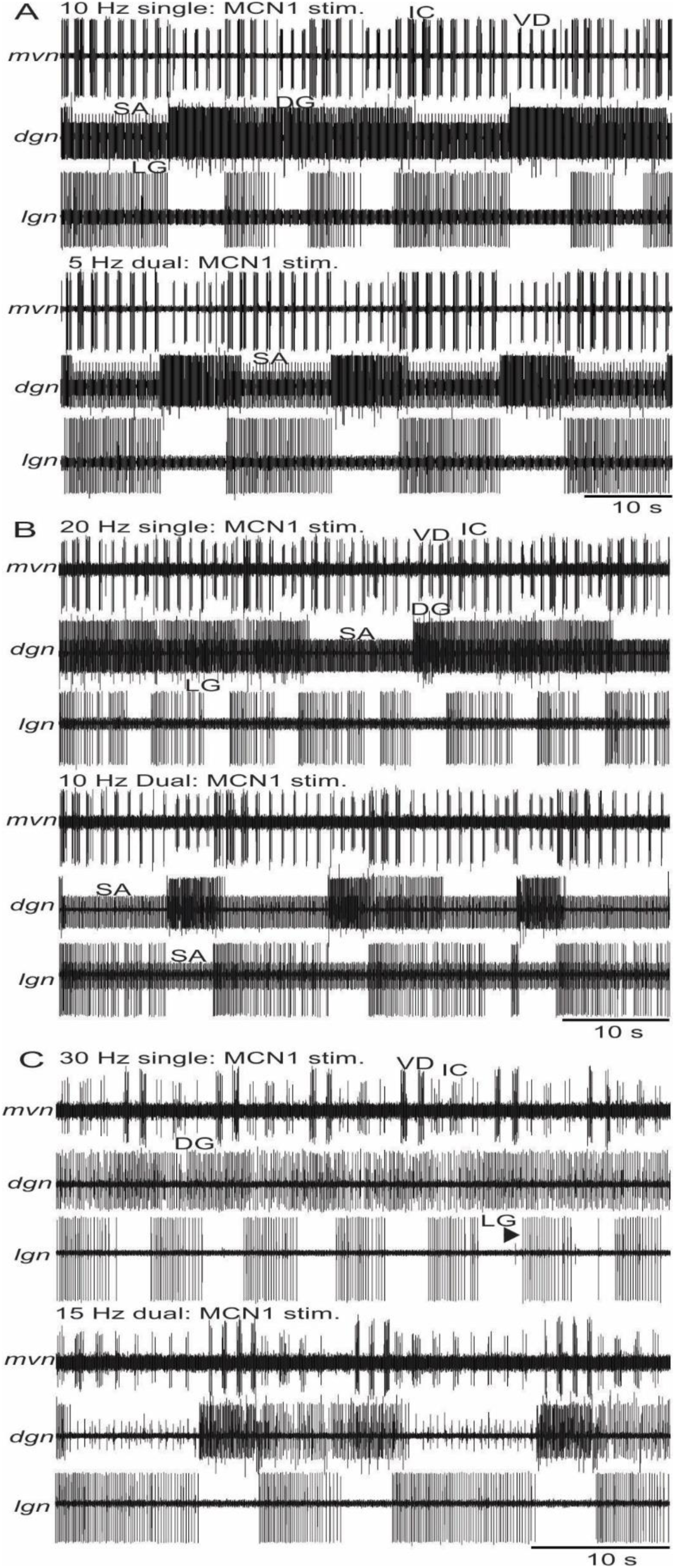
Example recordings of matched pairs of dual and single MCN1-driven gastric mill rhythms at an expanded time scale. Note that, unlike the poorly coordinated bursting between LG and DG, particularly during the single MCN1 stimulations, IC and VD neuron activity (*mvn*) consistently tracked LG activity. Each matched pair was recorded in the same experiment, but the different pairs come from different experiments. Matched MCN1 stimulations are compared at: (A) 5 Hz dual vs 10 Hz single; (B) 10 Hz dual vs. 20 Hz single, and (C) 15 Hz dual vs. 30 Hz single. Filtered: *mvn*-all recordings; *dgn*-10 Hz dual, 15 Hz dual, 30 Hz single.

The failure of DG activity to terminate during protraction was particularly pronounced during single MCN1 stimulations at 30 Hz. These stimulations never elicited LG/DG alternation in 75% or more of gastric mill rhythm cycles (n=0/16, 0%), and did so only sparingly (n=3/16; 19%) when threshold was reduced to 50% of cycles (Fig. 3). During the other 13 preparations (n=13/16; 81%), 30 Hz single MCN1 stimulation did not produce any cycles in which DG was silenced during protraction. This type of continuous DG activity throughout the gastric mill rhythm occurred during only two other MCN1 stimulation conditions (15 Hz dual stim., n=1/9, 11%; 20 Hz single stim., n=5/21, 24%).

### Distinct gastric mill rhythm parameters during matched dual and single MCN1 stimulations

There were also differences in several other gastric mill rhythm parameters between the matched dual and single MCN1 stimulations. For example, there was a longer gastric mill cycle period during the 5 Hz dual MCN1 stimulations than the 10 Hz single stimulations (n=14, p<0.001, one way RM-ANOVA, post-hoc Holm-Sidak test) (Figs. 4A, 5A). In contrast, the cycle period occurring during single MCN1_L_ (10 Hz) and MCN1_R_ (10 Hz) stimulation was comparable (n=14, p=0.98) (Fig. 5A). Despite the rhythm being slower during the dual 5 Hz stimulation, there was no difference in protraction duration between the dual and single stimulations (n=13, p=0.14, RM-ANOVA) (Fig. 5B). Instead, the longer cycle period resulted from an increased retraction duration (n=14, p<0.001, RM-ANOVA on Ranks, post-hoc Tukey test) (Fig. 5C). This selective prolonging of retraction during the dual 5 Hz MCN1 stimulations decreased the LG neuron duty cycle relative to the matched single stimulations (n=14, p<0.01; RM-ANOVA, post-hoc Holm-Sidak t-test) (Fig. 5D). The effects of 10 Hz_L_ and 10 Hz_R_ single MCN1 stimulations were indistinguishable for this and all other examined parameters (n=14, p>0.05) (Fig. 5).

**Figure 5.**
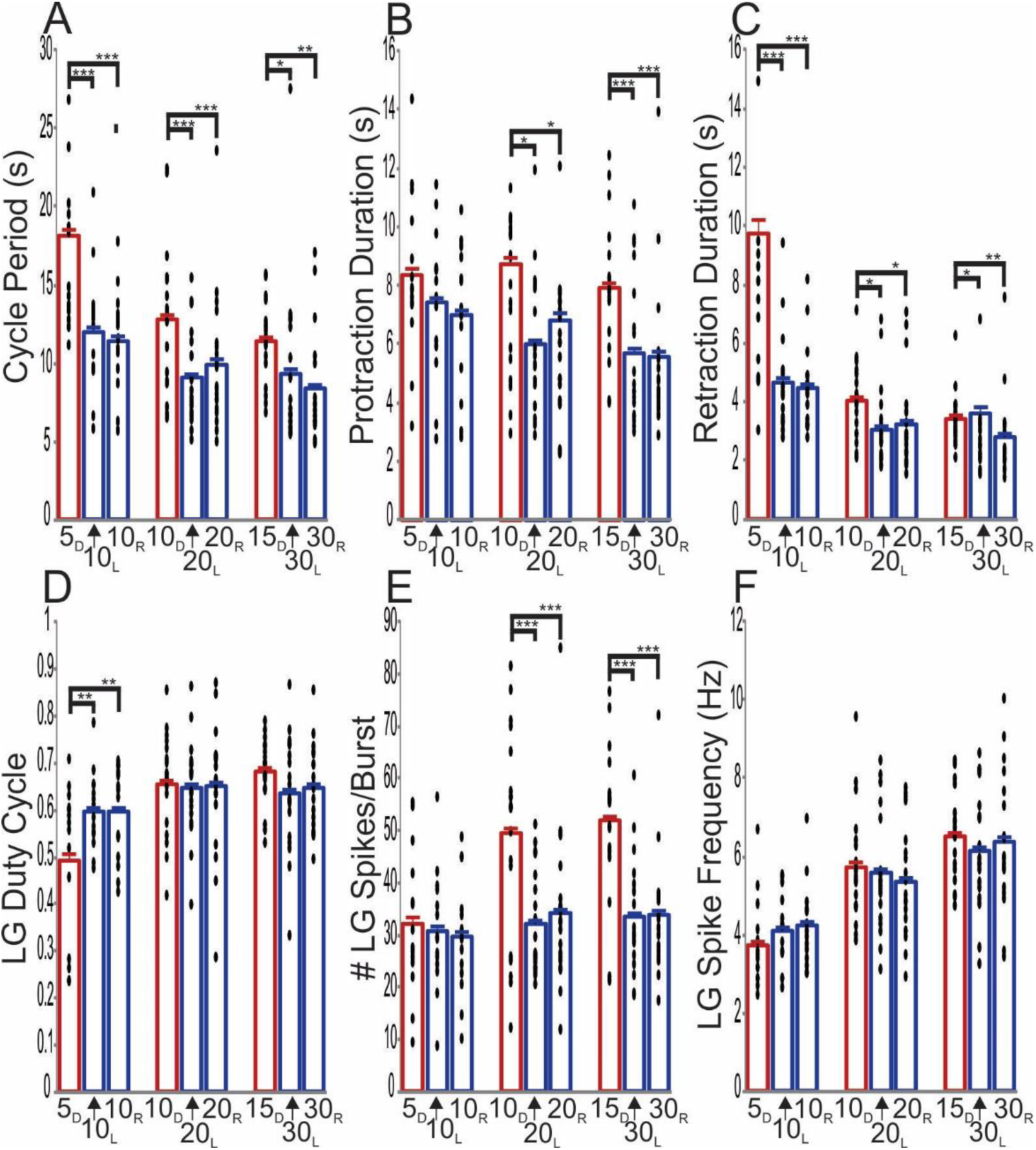
Gastric mill rhythm parameters are distinct during matched dual and single MCN1 stimulation. Bar graphs display the mean gastric mill rhythm values during three sets of firing rate- and pattern matched MCN1 stimulation protocols for (A) cycle period (outlier: 5 Hz Dual, 33.8 s), (B) protraction duration (outliers: 10 Hz Dual-19.8 s, 17.4 s; 20 Hz_R_-22.1 s), (C) retraction duration (outlier: 5 Hz Dual-26.2 s, 19.4 s; 30 Hz_L_, 18.5 s), (D) LG duty cycle, (E) number of LG spikes per burst, and (F) LG intraburst firing rate. The mean values for each experiment are shown within each bar (filled circles), excluding outliers which were omitted to optimize the y-axis dimensions. *p<0.05, **p<0.01, ***p<0.001; see text for n’s and statistical tests.

At the higher MCN1 stimulation frequency sets (10 Hz dual vs 20 Hz single, n=19; 15 Hz dual vs. 30 Hz single, n=18), protraction and retraction were both prolonged, and thus so was cycle period, during the dual MCN1 stimulations (Fig. 5A-C). In contrast, there was no distinction in these parameters between the single MCN1_L_ and MCN1_R_ stimulations at the same stimulation rate (Fig. 5A-C). The balanced increases in protraction and retraction duration during the higher dual stimulation rates resulted in there being no difference in the LG duty cycle between these conditions and their matched set of single MCN1 stimulations (Fig. 5C).

We also determined whether the LG neuron burst characteristics were differentially influenced by matched dual and single MCN1 stimulation. With respect to the number of LG spikes per burst, there was no difference at the lowest matched MCN1 stimulation rate (n=14, p=0.81, RM-ANOVA on Ranks) (Fig. 5E). In contrast, during both of the higher matched stimulation rates, the dual stimulations consistently elicited more LG spikes per burst than their matched single MCN1 stimulations (10 Hz dual vs. 20 Hz single: p<0.001, n=19; 15 Hz dual vs. 30 Hz single: p<0.001, n=18; RM-ANOVA, post-hoc Holm-Sidak test) (Fig. 5E). Although there were more LG spikes per burst during the two higher, dual MCN1 stimulation frequencies, there was no change in the LG intraburst firing rate within any of the three sets of stimulation conditions (Fig. 5F). Instead, these increased LG spike numbers reflected the prolonged LG burst duration (Fig. 5B).

Because both MCN1s are co-activated by all identified input pathways, we also compared the gastric mill rhythms when both MCN1s were co-stimulated (10 Hz each) vs. single MCN1 stimulation at the same frequency (n=13) (Fig. 6A,B). These stimulation conditions again elicited differences in some gastric mill rhythm parameters (Fig. 6B). For example, dual MCN1 stimulation (10 Hz ea.) elicited gastric mill rhythms exhibiting a longer cycle period (p=0.006) due to a prolonged protraction phase (p<0.001), with an increased LG duty cycle (p=0.003) containing more LG spikes per burst (p<0.001) generated at a higher firing rate (p<0.001) than during single MCN1 stimulation at 10 Hz (Fig. 6B). There was no difference in the retraction phase duration (p=0.455).

**Figure 6.**
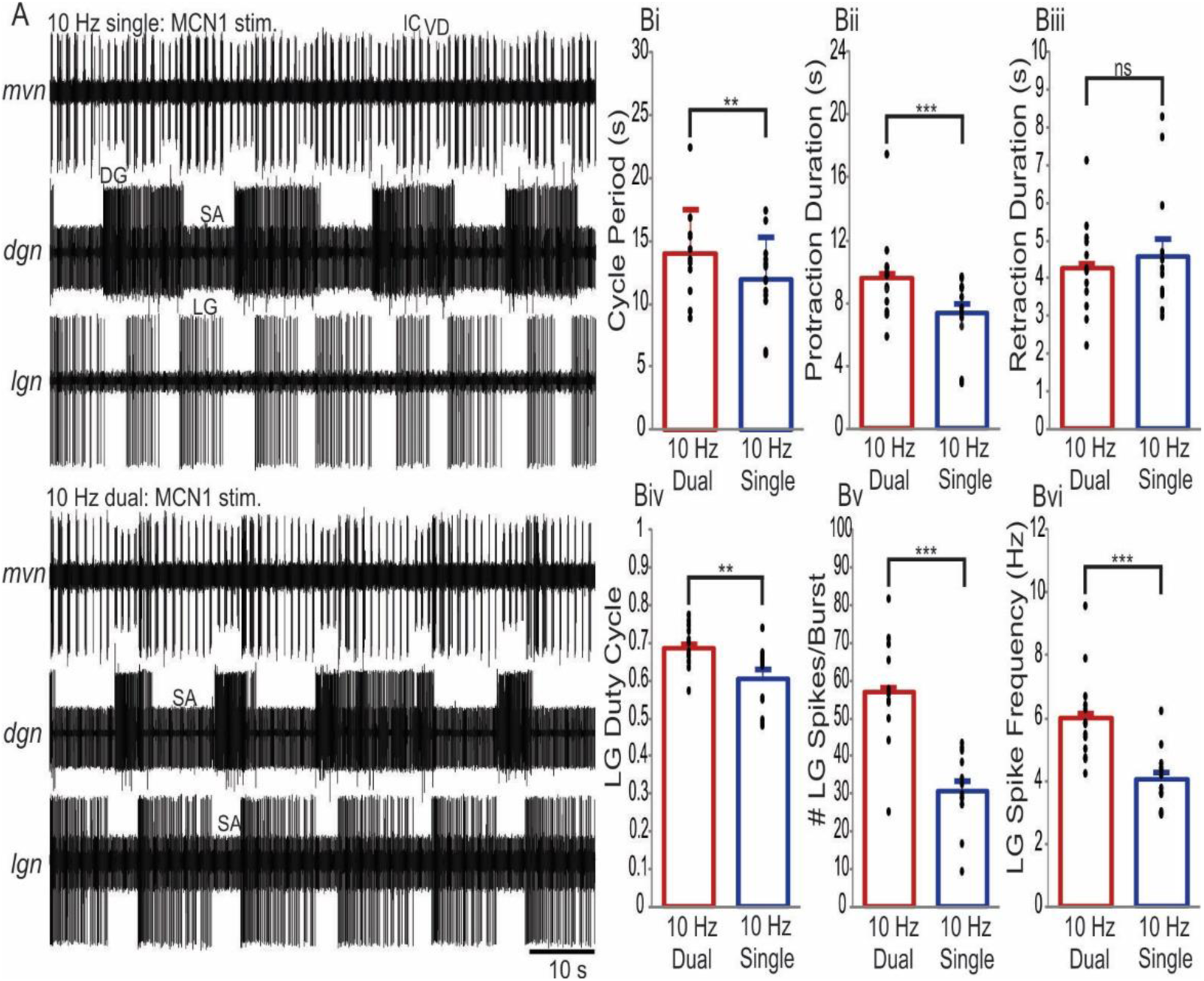
Dual MCN1 stimulation, at 10 Hz each, drives a protraction-prolonged gastric mill rhythm with a higher LG neuron intraburst firing frequency relative to single MCN1 stimulation at 10 Hz. **(A)** Example gastric mill rhythms driven by dual MCN1 stimulation (10 Hz ea.) and single MCN1 stimulation at 10 Hz. Filtered: 10 Hz single-*mvn*; 10 Hz dual-*mvn, dgn*. **(B)** Bar graphs compare the mean value for different gastric mill rhythm parameters during dual-MCN1 stimulation (10 Hz ea.) and single MCN1 stimulation (10 Hz; n=13). Filled circles spanning each bar represent the mean values from each experiment. (Bi) Cycle period: p=0.006. (Bii) Protraction duration: p<0.001. (Biii) Retraction duration: p=0.455. (Biv) LG duty cycle: p=0.003. (Bv) #LG spikes/burst: p<0.001. (Bvi) LG Intraburst firing frequency: p<0.001. Bi, Bii, Bii, Bvi: Wilcoxon Signed Rank Test; BiV, Bv: Paired t-Test.

In addition to differences in the mean value of some gastric mill rhythm parameters between matched dual and single MCN1 stimulation frequencies, we assessed parameter variability across experiments in relation to the mean using the Coefficient of Variation (CV). Statistically significant differences occurred primarily at the lowest stimulation rate (5 Hz dual vs. 10 Hz single). For example, the CV across experiments for cycle period was higher for the 10 Hz single MCN1 stimulations (dual MCN1 vs: MCN1_L_, p=0.028; MCN1_R_, p=0.031; RM-ANOVA, Holm-Sidak post-hoc test; n=14) (Fig. 7A). Some of the studied parameters whose mean values were not different nevertheless exhibited differences in the CV, as was the case for the protraction duration and number of LG spikes per burst (Figs. 5,7). For example, the protraction duration CV during the single 10 Hz stimulations for MCN1_L_ was 0.27 ± 0.01, while it was 0.17 ± 0.01 for the dual 5 Hz stimulation (n=14; p=0.045, RM-ANOVA, Holm-Sidak post-hoc test). However, despite the 10 Hz stimulation of MCN1_R_ having the same protraction duration CV as MCN1_L_ (0.27 ± 0.01), this value was not different from that of the matched 5 Hz dual stimulations (p=0.062, RM-ANOVA, Holm-Sidak post-hoc test). With respect to the number of LG spikes per burst, single MCN1 stimulations at 10 Hz produced a higher CV across experiments than the dual 5 Hz stimulation (dual MCN1 vs: MCN1_L_, p=0.03; MCN1_R_, p=0.042; RM-ANOVA, Holm-Sidak post-hoc test; n=14). In contrast, the CV for retraction duration, LG duty cycle and LG firing frequency was comparable for the matched dual (5 Hz ea.) and single (10 Hz) MCN1 stimulations (Fig. 7C,D,F). Few parameters exhibited CV differences between the higher frequency-matched MCN1 stimulations, and in no cases did both single stimulations differ from the matched dual stimulation (Fig. 7).

**Figure 7.**
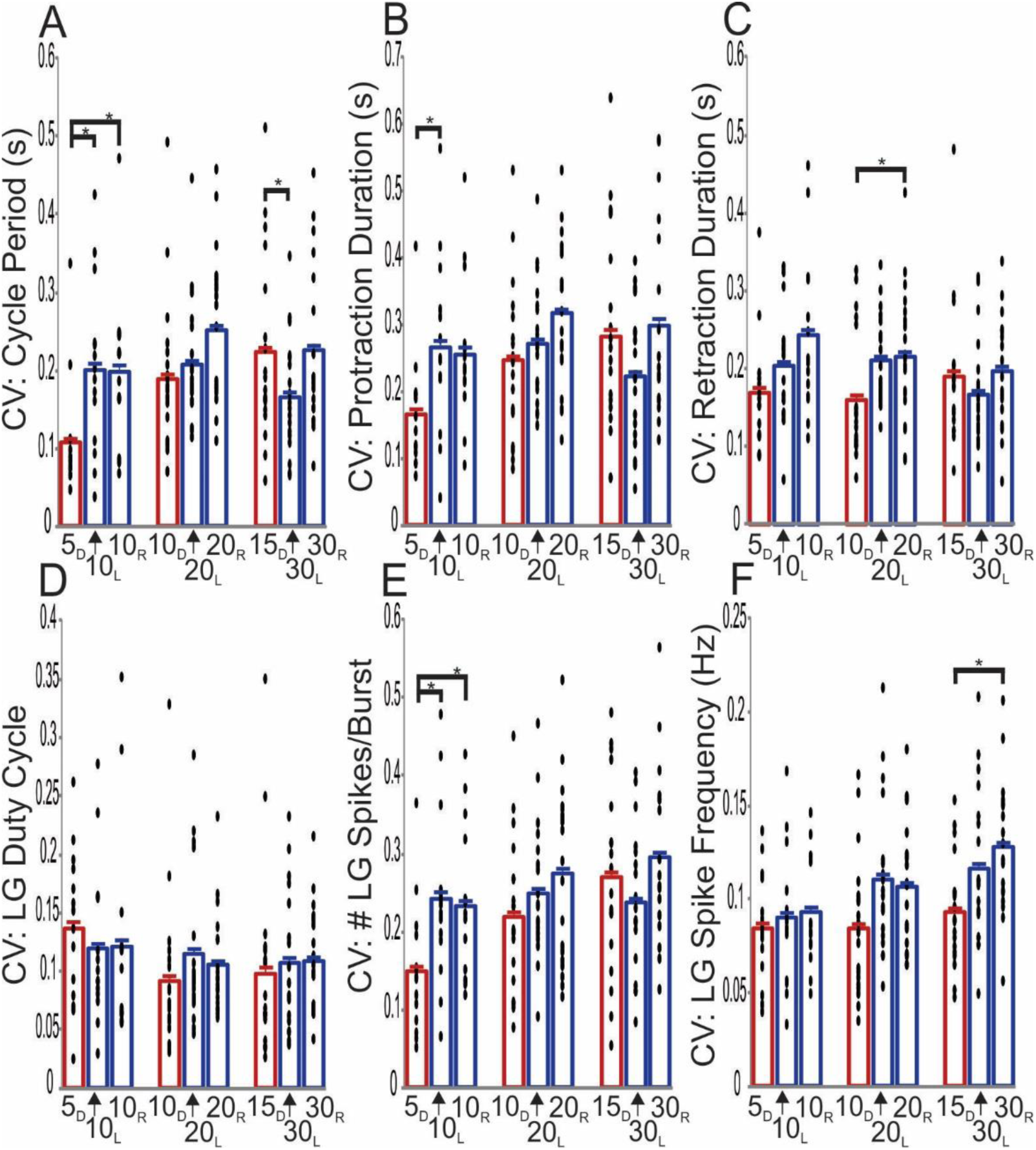
The inter-experiment coefficient of variability (CV) for some gastric mill rhythm parameters is distinct between matched dual- and single MCN1 stimulation. Bar graphs display the mean gastric mill rhythm CV values during three sets of firing rate- and pattern matched MCN1 stimulation protocols for (A) cycle period, (B) protraction duration, (C) retraction duration, (D) LG duty cycle, (E) number of LG spikes per burst, and (F) LG intraburst firing rate. The mean values for each experiment are shown within each bar (filled circles). *p<0.05; RM-ANOVA plus Holm-Sidak post-hoc test for all comparisons except cycle period for 15 Hz dual vs. 30 Hz single (RM-ANOVA on Ranks plus Tukey t-test). See text for n’s and statistical test values.

### The LG neuron bursts weaken but do not eliminate MCN1_STG_ synaptic actions

The fact that DG neuron activity was not always limited to the retraction phase in these experiments, but exhibited a reduced firing rate when active during protraction (Figs. 2-4), suggested that the LG firing frequency during these rhythms weakened but did not eliminate MCN1_STG_ transmitter release. We tested this hypothesis by comparing the influence of natural and strengthened LG bursts on (a) DG neuron activity, and (b) the pyloric cycle period, during gastric mill rhythms driven by matched dual (10 Hz) and single (20 Hz) MCN1 stimulation. In these experiments we strengthened some LG bursts by increasing its within-burst firing rate via intracellular depolarizing current pulses (see Methods). The pyloric cycle period is a useful assay because MCN1 directly excites the pyloric rhythm, reducing the pyloric cycle period, whereas LG activity only influences this rhythm (i.e. increases the pyloric cycle period) via its inhibition of MCN1_STG_ (Bartos and Nusbaum, 1997).

When LG neuron activity was strengthened during matched dual (10 Hz) and single (20 Hz) MCN1 stimulations, its increased firing rate was not different across protocols (20 Hz_L_: 13.9 ± 0.4 Hz; 20 Hz_R_: 13.4 ± 0.4 Hz; 10 Hz_Both_: 13.1 ± 0.6 Hz; n= 5 exps, 10 LG dep./condition; p= 0.111, RM-ANOVA). These increased firing rates were consistently higher than during the control LG bursts (p<0.001 for each protocol, RM-ANOVA, Holm-Sidak post-hoc test), which ranged from 4.5 Hz – 6.0 Hz.

For gastric mill rhythm cycles where DG neuron activity extended through the time of the LG burst, enhanced LG activity eliminated this prolonged DG activity in 10/20 cycles during 20 Hz single MCN1 stimulations (50%; n=5 exps, 2 LG current injections per MCN1 stimulation). In the remaining 10 cycles, the DG firing rate was weakened but not terminated (e.g. Fig. 8). This extended DG activity was more consistently eliminated during the dual 10 Hz MCN1 stimulations (9/10 cycles 90%; n=5 exps, 2 LG current injections/MCN1 stim.; p=0.049, Fisher’s Exact test), supporting the suggestion that LG more effectively regulated MCN1 activity when MCN1 fired at a lower rate (Fig. 8).

**Figure 8.**
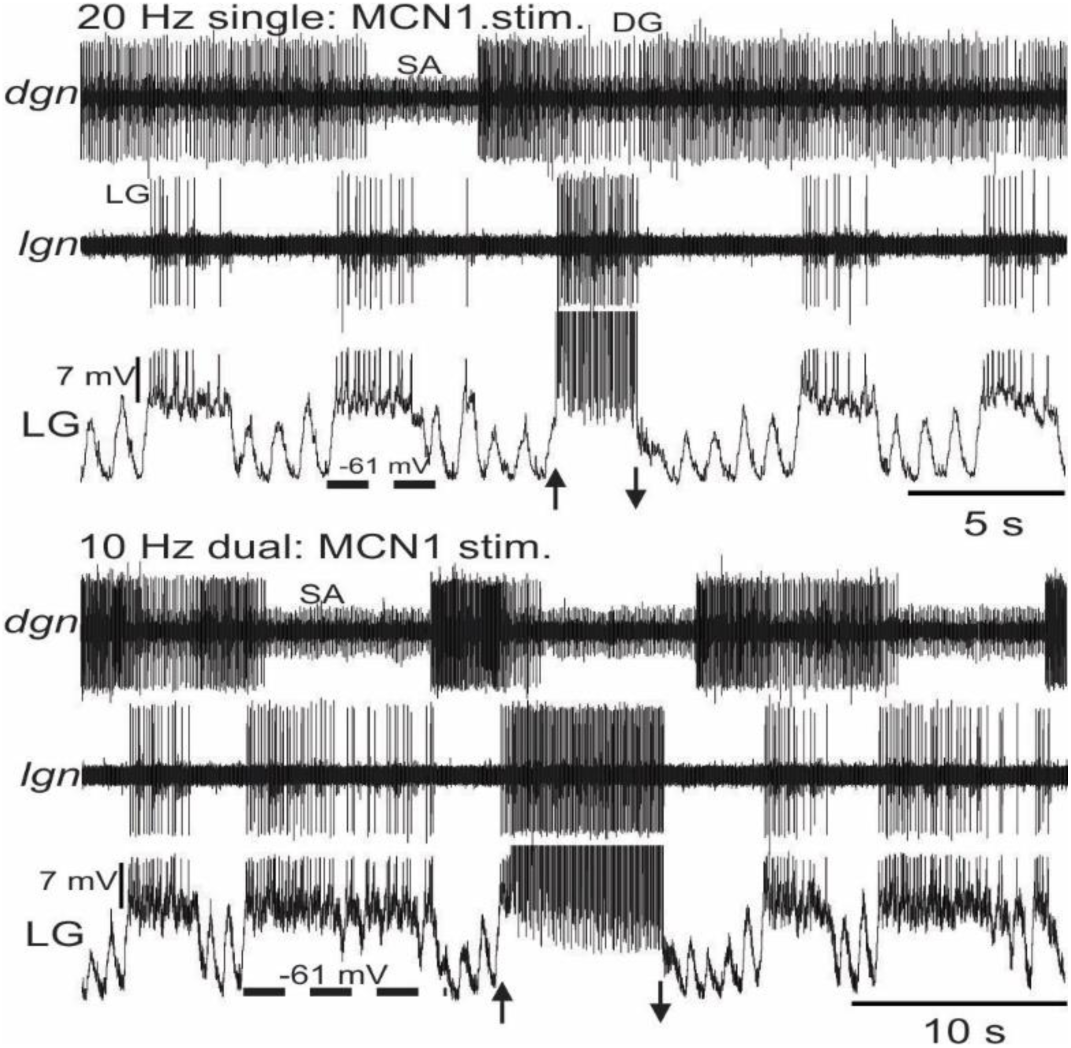
Increasing the LG neuron intraburst firing frequency more effectively regulates DG neuron burst termination during the MCN1-gastric mill rhythm. Example recordings show the effect of increasing the LG neuron firing rate on the preceding/overlapping DG neuron burst during gastric mill rhythms driven by matched dual (10 Hz) and single (20 Hz) MCN1 stimulation in the same experiment. In both recordings, most of the bracketing control cycles show DG activity persisting through the LG burst, albeit at a lower firing rate than before and after the LG burst. The LG firing rate is more evident in the extracellular recording (*lgn*) due to the current injection pulses present in the intracellular recording (LG). Up and down arrows indicate onset and termination, respectively, of each set of depolarizing current pulses.

We evaluated the impact of an increased LG firing frequency on the pyloric cycle period by first normalizing the pyloric cycle period during strengthened LG firing to that during the preceding, control LG burst (‘LG Dep. ratio’; see Methods). We then compared this normalized value to a control value obtained by dividing the pyloric cycle period during the same, preceding control LG burst by the cycle period during the control LG burst two prior to the depolarized LG burst (‘Control ratio’). The control pyloric cycle period ratios were comparable across all MCN1 stimulation protocols (n=10 per protocol, p=0.407, RM-ANOVA on Ranks).

During all three sets of MCN1 stimulation, the LG Dep. ratio was larger than the Control ratio (20 Hz_L_: LG dep. ratio, 1.11 ± 0.03; Control ratio, 1.00 ± 0.03, p=0.004; 20 Hz_R_: LG dep. ratio, 1.18 ± 0.06; Control ratio, 0.99 ± 0.01, p=0.004; 10 Hz_Both_: LG dep. ratio, 1.07 ± 0.01; Control ratio, 0.99 ± 0.01, p=0.001; n=5, Signed Rank Test [20 Hz_L,R_], Paired t-test [10Hz_Both_]) (Fig. 9). The increased ratios that occurred during the LG current injections indicated that the pyloric cycle period at these times was prolonged relative to that during the control LG bursts. Consistent with the limited LG/DG alternation reported above (Figs. 2-4), and the impact of an increased LG firing rate on the timing of DG neuron bursts, these pyloric rhythm results suggested that the strengthened LG bursts more strongly inhibit MCN1_STG_ transmitter release.

**Figure 9.**
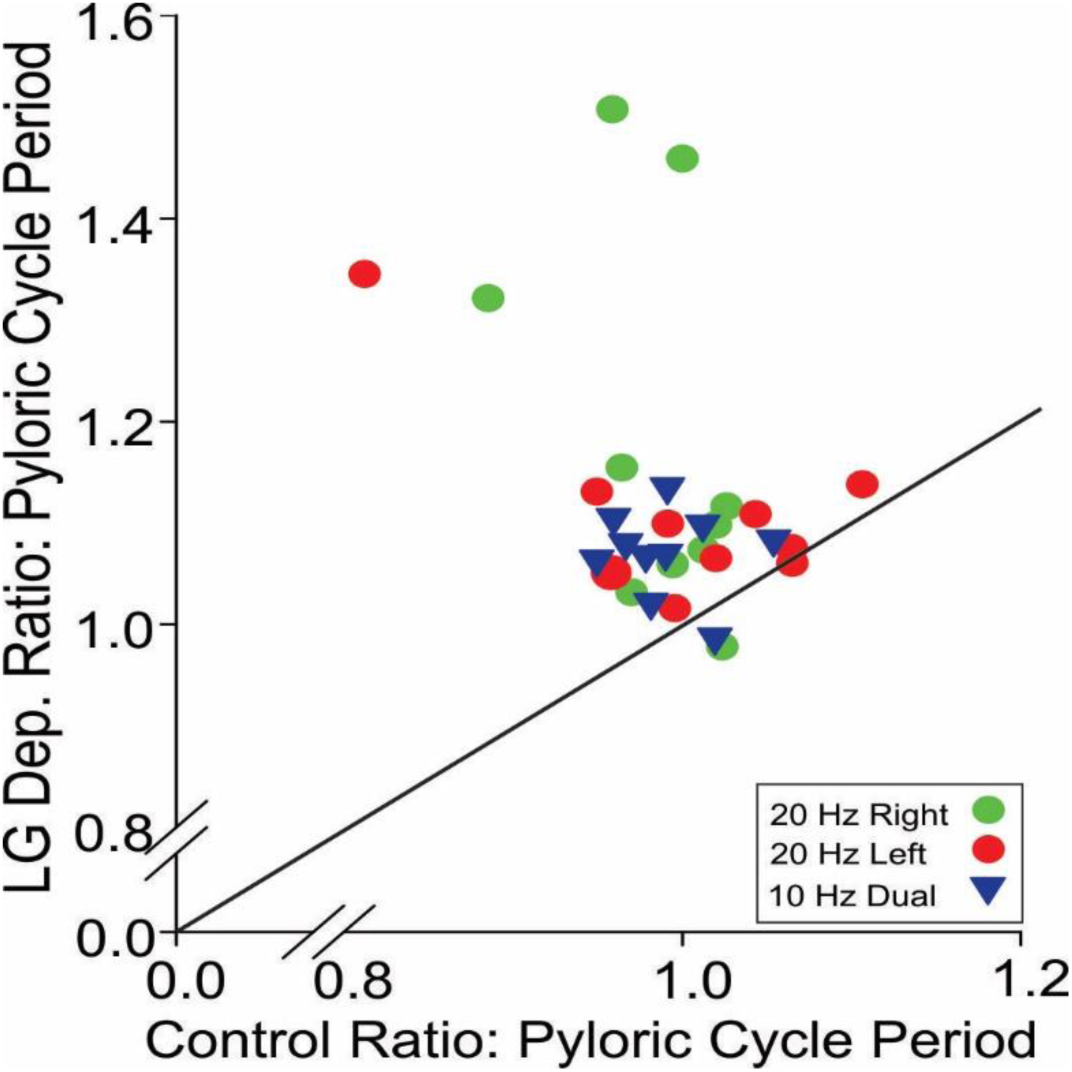
The pyloric cycle period is prolonged by increasing the intraburst LG firing frequency. Scatter plot showing the ratio of the pyloric cycle period during an enhanced LG burst to that of the preceding LG burst (LG Dep. Ratio) relative to its control ratio, during gastric mill rhythms driven by single (20 Hz) or dual (10 Hz) MCN1 stimulation (n=10: 5 experiments, 2 LG Dep. per experiment for each single and dual MCN1 stimulation). Diagonal line slope = 1.

### Metabotropic activation of MCN1 mimics the effects of extracellular MCN1 stimulation

We used *ion* stimulation to selectively drive MCN1 because the only other CoG neuron (MCN5) that projects to the STG through the *ion* has a smaller diameter axon, enabling selective activation of MCN1 by extracellular stimulation (Coleman et al., 1992; Coleman and Nusbaum, 1994; Norris et al., 1996). We nevertheless considered the possibility that some MCN5 stimulation contributed to our results by comparing the influence of *ion* stimulation with that resulting from stimulating the *dpon*, after bisecting the *son* medial to the *dpon* (see Methods) (Fig. 1A). This approach stimulates a mechanosensory pathway (VCN neurons) which triggers a long-lasting activation of MCN1, but not MCN5 (Beenhakker et al., 2004).

The MCN1 firing rate resulting from *dpon* stimulation in different experiments ranged from 9.5 Hz – 26 Hz. To match the *ion* and *dpon* activation of MCN1 in each experiment, we first triggered a VCN-activation of MCN1 and recorded the resulting gastric mill rhythm. In parallel, we determined the VCN-triggered MCN1 firing rate and used that same rate to subsequently stimulate the *ion*. We then followed each *ion* stimulation with a second *dpon* stimulation. Little or no MCN5 activity was evident in the *ion* recordings during the VCN-triggered gastric mill rhythms. We assayed the gastric mill rhythm response by analyzing the same gastric mill rhythm parameters as above (Fig. 5) and determining the gastric mill rhythm-timed activity of the DG neuron.

The gastric mill circuit response to *dpon* and *ion* stimulation was comparable for five of the six analyzed parameters, including cycle period, protraction- and retraction duration, number of LG spikes per burst, and the LG duty cycle (n=5) (Fig. 10). The only analyzed parameter with distinct values was the LG neuron firing frequency (*dpon* stim. 1: 6.6 ± 1.1 Hz; *ion* stim.: 5.9 ± 0.9 Hz; *dpon* stim. 2: 6.5 ± 1.0 Hz; n=5; *dpon*1 vs. *ion*: p=0.03; *dpon*2 vs. *ion*: p= 0.04; *dpon*1 vs *dpon* 2: p=0.67, RM-ANOVA, Holm-Sidak post-hoc test) (Fig. 10F). Similarly, the matched MCN1 firing rates from *dpon* and *ion* stimulation resulted in comparable DG neuron activity patterns. In none of these experiments was DG activity limited to the retraction phase, nor were there any gastric mill rhythms in which the LG and DG bursts alternated in at least 50% of the cycles (n=5).

**Figure 10:**
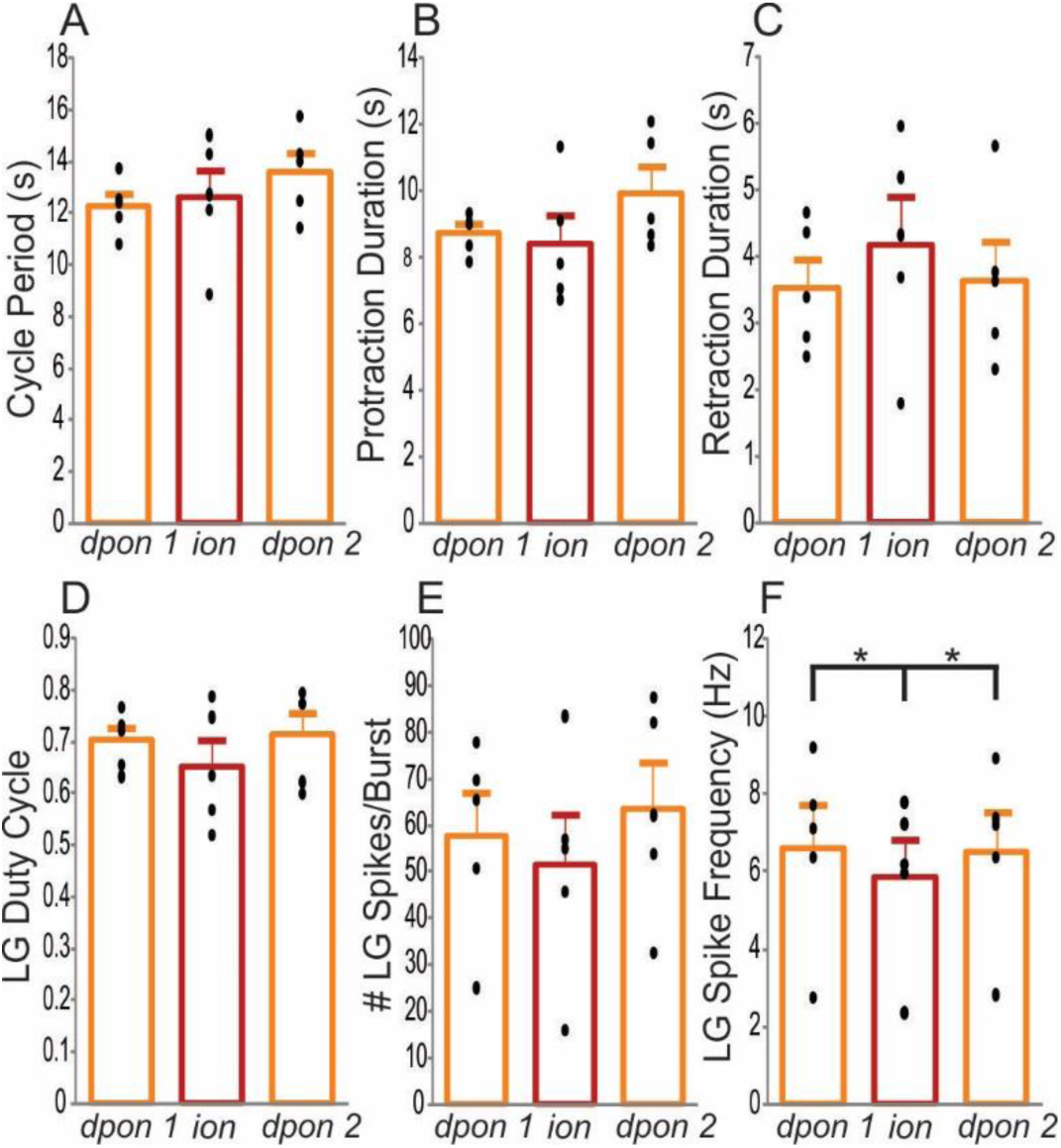
VCN stimulation with the *sons* bisected elicits a MCN1-gastric mill rhythm comparable to that resulting from *ion* stimulation using the same MCN1 firing rate. Bar graphs display the mean gastric mill rhythm values triggered by VCN stimulation, before (*dpon1*) and after (*dpon2*) MCN1 (*ion*) stimulation for (A) cycle period, (B) protraction duration, (C) retraction duration, (D) LG duty cycle, (E) #LG spikes per burst, and (F) LG intraburst firing rate. Mean values for each experiment are shown within each bar (filled circles; n=5). *p=0.04, RM-ANOVA plus Holm-Sidak post-hoc test.

## DISCUSSION

In this paper we determined that firing rate- and pattern matched dual and single stimulation of the paired projection neuron MCN1 does not elicit equivalent gastric mill motor patterns. These different outcomes appear due, at least partly, to a mismatch in the firing rate of MCN1 and the gastric mill circuit neuron LG, which regulates MCN1 transmission in the STG via presynaptic inhibition (Coleman and Nusbaum, 1994; Coleman et al., 1995). Specifically, the dual MCN1 stimulations more consistently elicited fully coordinated gastric mill cycles. This coordination distinction was particularly prevalent at the lowest MCN1 stimulation frequency comparison, where the gastric mill rhythm was fully coordinated during every cycle in 80% of the 5 Hz dual stimulations but in none of the matched 10 Hz single stimulations. These low frequency dual stimulations also exhibited lower variability in several parameters across experiments. This outcome suggests that acute loss of one MCN1 would compromise behavioral performance to the extent that effective chewing involves coordinated rhythmic protraction and retraction of the teeth (Heinzel et al., 1993; Diehl et al., 2013). The functional consequences after such a loss might be improved over longer durations, however, given that long-term compensation does occur in at least some rhythmic motor systems (Büschges et al., 1992; Sánchez et al., 2000; Sakurai and Katz, 2009; Fink and Cafferty, 2016; Sakurai et al., 2016; Brown and Martinez, 2018; Puhl et al., 2018).

Insofar as most gastric mill circuit neurons are also the motor neurons for the system, the other rhythm parameter differences that we identified suggest differences would also occur in the response dynamics of at least some gastric mill muscles, producing changes in the timing and/or strength of teeth movements (Stein et al., 2006; Diehl et al., 2013). Whether these latter changes would compromise chewing behavior, however, remains to be determined. In all comparisons, there were no differences between the matched single stimulations (MCN1_L_ vs. MCN1_R_), which is in agreement with previous evidence suggesting that the two MCN1s are functionally equivalent. There do not appear to be other studies that have performed such a comparison on a projection neuron population.

The role(s) of presynaptic regulation of projection- and sensory neuron inputs to neuronal circuits remains under-explored, despite its prevalence in many neural systems (Nusbaum, 1994; Coleman and Nusbaum, 1994; Sillar and Simmers, 1994; Krieger et al., 1996; Cochilla and Alford, 1999; Westberg et al., 2000; Takahashi and Alford, 2002; Evans et al., 2003; Hurwitz et al., 2005; Barrière et al., 2008; Blitz and Nusbaum, 2008, 2012; Jing et al., 2011; Wang, 2012; McGann, 2013; Sirois et al., 2013; Blitz et al., 2019). For the MCN1-gastric mill rhythm, the pivotal role of LG neuron presynaptic inhibition of MCN1_STG_ supports the hypothesis that this circuit design favors dual MCN1 activity over firing rate-matched single MCN1 activity. This hypothesis is consistent with the greater effectiveness for this presynaptic action during the dual MCN1 stimulations, where each MCN1 fired at half the rate of the matched single MCN1 stimulations, and by our finding that the LG firing rate was equivalent within each matched set of MCN1 stimulations. This outcome resulted from the LG synaptic action reducing but not eliminating MCN1 transmitter release in these experiments, as evidenced by the observations that increasing the LG intraburst firing rate during both dual and single MCN1 stimulation more consistently limited DG neuron activity to the retraction phase, and further weakened the pyloric rhythm. LG only regulates these two events via its inhibitory action on MCN1_STG_ (Coleman and Nusbaum, 1994; Bartos and Nusbaum, 1997). This is the first indication in the biological system that the gastric mill rhythm-timed LG bursts reduce but do not eliminate MCN1_STG_ transmitter release, although a previous computational model did support this possibility (DeLong et al., 2009a).

The prolonged gastric mill cycle period during the dual MCN1 stimulations resulted from selectively prolonged retraction during dual 5 Hz MCN1 stimulation, whereas both phases were prolonged during the dual 10 Hz and 15 Hz MCN1 stimulations. This distinction suggests that different cellular/synaptic mechanisms underlie cycle period regulation during different MCN1 firing rates. Selectively prolonged retraction occurs when the rate of I_MI_ accumulation in LG is reduced at times when there is little or no change in the strength of Int1 inhibition (Beenhakker et al., 2005; DeLong et al., 2009a,b). In the present study, this event may have resulted from a relatively low release rate of CabTRP Ia peptide when each MCN1 is firing at 5 Hz, a firing rate which commonly generates a near-threshold level of neuropeptide release (Vilim et al., 1996, 2000; Liu et al., 2011; Ding et al., 2019). MCN1-released CabTRP la peptide activates I_MI_ in LG, and 5 Hz is the approximate threshold firing frequency for the MCN1-driven gastric mill rhythm (Wood et al., 2000; Kirby and Nusbaum, 2007; DeLong et al., 2009a). Selectively prolonged retraction may also result from I_MI_ declining to a lower level at the end of each LG burst during the 5 Hz dual MCN1 stimulations relative to the matched 10 Hz single stimulations, due to LG more strongly inhibiting MCN1_STG_ transmitter release at the lower MCN1 firing rate. After a stronger LG inhibition of MCN1_STG_, more time would be needed for sufficient I_MI_ to build-up and trigger the next LG burst (DeLong et al., 2009a).

How the dual 10 Hz and 15 Hz MCN1 stimulations prolonged both gastric mill rhythm phases relative to their matched single stimulations is less clear. However, MCN1 activity does prolong both phases when Int1 inhibition of LG is strengthened to a greater degree than the parallel rate of I_MI_ buildup (Beenhakker et al. 2005). At the higher MCN1 stimulation frequencies the dual stimulations may elicit an altered balance of CabTRP Ia and GABA release, relative to their matched single stimulations, such that the MCN1 GABAergic excitation of Int1 becomes relatively more strengthened than the peptidergic excitation of LG. Neuropeptide and small molecule transmitter release can have different Ca^2+^ and/or firing rate-dependencies (Whim and Lloyd, 1989; Peng and Zucker, 1993; Liu et al., 2011; Nusbaum et al., 2017). Also, as above, the more effective LG inhibition of MCN1_STG_ during the dual stimulations would likely more completely reduce I_MI_ amplitude at the end of each LG burst, contributing to a longer duration for I_MI_ buildup during each subsequent retraction phase.

In contrast to these experiments, where optimal generation of a fully coordinated gastric mill rhythm resulted from dual MCN1 stimulation at 5 Hz, the MCN1 firing rate is often considerably higher (25-30 Hz) during fully coordinated gastric mill rhythms triggered by the VCN- or POC neurons in the complete STNS (Beenhakker and Nusbaum, 2004; Blitz and Nusbaum, 2012). However, the VCN- and POC-triggered gastric mill rhythms are driven by co-activating MCN1 and CPN2, another CoG projection neuron. Also, during these latter rhythms the firing pattern of both projection neurons is not tonic but is rhythmically coordinated with the gastric mill- and pyloric rhythms, due to synaptic input from the gastric mill- and pyloric circuit interneurons Int1 and AB. Circuit-linked activity patterns in projection neurons that drive rhythmic motor patterns are common across systems (Weeks and Kristan, 1978; Rosen et al., 1991; Norris et al., 1994, 1996; Puhl et al., 2012; Grillner and El Manira, 2020). It remains to be determined whether some or all of these distinctions, as well as the rhythmic sensory feedback that would be present in vivo, compensate for the degraded coordinated activity that occurred under many of our experimental conditions in the isolated STG. Although rhythmic sensory feedback is not necessary for core rhythm generation in most motor systems, such feedback commonly sculpts the final motor pattern in vivo (Wolf and Pearson, 1988; Hooper et al., 1990; Büschges et al., 1992; Combes et al., 1999; Eisenhart et al., 2000; Marder and Bucher, 2001; Smarandache and Stein, 2007; Hedrich et al., 2009; Buschges et al., 2011; Akay et al., 2014). Additionally, regarding this latter issue, DG neuron-mediated muscle contraction activates the muscle stretch-sensitive neuron GPR (Katz et al., 1989). GPR activity, in turn, excites MCN1 and CPN2 in both CoGs, influences the gastric mill rhythm generator neurons in the STG, and can entrain the gastric mill rhythm (Blitz et al., 2004; Beenhakker et al., 2005). Thus, this sensory feedback pathway may normally provide a correction signal that maintains MCN1-driven gastric mill rhythm coordination in vivo.

The superior coordination of gastric mill rhythms driven by dual MCN1 stimulation provides a reasonable explanation for why all identified inputs to MCN1 influence both copies, despite their presence in separate ganglia (Beenhakker et al., 2004, 2005; Blitz et al., 2004, 2008; Christie et al., 2004; Hedrich et al., 2009; Blitz and Nusbaum, 2012; White et al., 2017). This system design also provides a cautionary note for efforts focused on enabling recovery of function after partial loss of pathways that drive behavior. Increasing the firing rate of the remaining population in a compromised pathway might be expected to compensate for partial loss of a population, but this is not the only possible outcome. For example, as noted above, neurons often exhibit firing rate-dependent changes in neurotransmitter release that can alter their ionotropic and metabotropic actions, leading to qualitative changes in circuit output. Additionally, insofar as neurotransmitter release for many neurons is also regulated locally at axon terminals, the balance between the projection neuron firing rate and that of the regulating presynaptic neuron can be pivotal to maintaining a coordinated motor pattern, as is the case for MCN1. In conclusion, the gastric mill microcircuit is more effectively driven by co-activating the paired projection neuron MCN1 than by its firing rate- and pattern matched single stimulation, at least in the isolated nervous system.

## Acknowledgements

We thank Mr. Logan Fickling for helpful feedback on earlier versions of the manuscript.

## List of Abbreviations

AB: anterior burster (neuron)
AGR: anterior gastric receptor (neuron)
AM: anterior median (neuron)
*C. borealis*: *Cancer borealis*
CabTRP la: *Cancer borealis* tachykinin-related peptide la
CoG: commissural ganglion
CV: coefficient of variation
DCC: discontinuous current clamp
DG: dorsal gastric (neuron)
*dgn*: dorsal gastric nerve
*dpon*: dorsal posterior oesophageal nerve
*dvn*: dorsal ventricular nerve
eEPSP: electrical excitatory postsynaptic potential
GABA: gamma amino butyric acid
GM: gastric mill (neuron)
g_MI_: modulator-activated voltage-dependent inward conductance
i: ionotropic (transmission)
IC: inferior cardiac (neuron)
I_MI_: <u>m</u>odulator-activated inward current
Int1: interneuron 1
*ion*: inferior oesophageal nerve
LG: lateral gastric (neuron)
*lgn*: lateral gastric nerve
*lvn*: lateral ventricular nerve
m: metabotropic (transmission)
MCN1: modulatory commissural neuron 1
MCN1_L_: left MCN1
MCN1_R_: right MCN1
MCN1_STG_: STG terminals of MCN1
MCN5: modulatory commissural neuron 5
MG: medial gastric (neuron)
*mvn*: medial ventricular nerve
OG: oesophageal ganglion
PD: pyloric dilator (neuron)
*pdn*: pyloric dilator nerve
Pro: protraction
Ret: retraction
RM-ANOVA: repeated measures-analysis of variance
SA: stimulus artifact
*son*: superior oesophageal nerve
STG: stomatogastric ganglion
stn: stomatogastric nerve
STNS: stomatogastric nervous system
VCNs: ventral cardiac neurons
VD: ventricular dilator (neuron)

